# Decreased transcription factor binding levels nearby primate pseudogenes suggests regulatory degeneration

**DOI:** 10.1101/024026

**Authors:** Gavin M. Douglas, Michael D. Wilson, Alan M. Moses

## Abstract

Characteristics of pseudogene degeneration at the coding level are well-known, such as a shift towards neutral rates of nonsynonymous substitutions and gain of frameshift mutations. In contrast, degeneration of pseudogene transcriptional regulation is not well understood. Here, we test two predictions of regulatory degeneration along the pseudogenized lineage: (1) decreased transcription factor binding and (2) accelerated evolution in putative cis-regulatory regions.

We find evidence for decreased TF binding levels nearby two primate pseudogenes compared to functional liver genes. We also find evidence for pseudogene-lineage-specific relaxation of sequence constraint on a fragment of the promoter of the primate pseudogene urate oxidase (*Uox*) and a nearby cis-regulatory module (CRM). However, the majority of TF-bound sequences nearby pseudogenes do not show evidence for lineage-specific accelerated rates of evolution. We conclude that decreases in TF binding level could be a marker for regulatory degeneration, while sequence degeneration in most CRMs may be obscured by background rates of TF binding site turnover.

---

Regulatory evolution is thought to be a key mechanism underlying the diversity of closely related species (Britten and Davidson 1969; King and Wilson 1975). For example, deletions of conserved regulatory elements have been suggested to underlie human-specific traits (McLean et al. 2011). Despite the importance of regulatory degeneration for human evolution, little is known in general about how regulation evolves at the molecular level once selective constraint has been lifted. By understanding how transcriptional regulation degenerates, we can help resolve which aspects of transcriptional regulation natural selection is preserving. Here, we study transcription factor binding and cis-regulatory module (CRM) evolution near human pseudogenes to characterize the process of regulatory degeneration, which we refer to as “pseudo-enhancerization” in analogy with pseudogenization (the process whereby natural selection fails to preserve genes). We focus on regulatory degeneration near unitary pseudogenes, because these represent true losses of function from a species rather than degeneration of a redundant copy.

Two classic examples of unitary pseudogenes in human are urate oxidase, *Uox*, and L-gulonolactone oxidase, *Gulo*, which are functional in the livers of most mammals. *Uox* removes excess nitrogen from the body by metabolizing uric acid (Gustafsson and Unwin 2013) and was pseudogenized independently in both human and gibbon (Zhang et al. 2010). *Gulo* is responsible for encoding the protein required for the final step in the vitamin C (ascorbic acid) synthesis pathway and was lost in the ancestor of human and macaque (Drouin et al. 2011).

There are a number characteristics of pseudogenization that are used to identify pseudogenes and are all present in *Uox* and *Gulo* (Zhang et al. 2010). Firstly, the former coding sequence of pseudogenes evolves faster than the orthologous coding regions in species where the loci are still functional. Similarly, pseudogenes exhibit a K_a_/K_s_ (ratio of nonsynonymous to synonymous substitution rates) that approaches 1 as evolutionary distance increases. Both of these characteristics are due to relaxed purifying selection acting at the sequence level.

In contrast, it is unknown whether regulatory sequences show similar signatures of relaxed selection. Regulatory elements that regulated the functional ancestors of pseudogenes are expected to degrade following the pseudogenization event. In fact, it is possible that the first steps of pseudogenization actually occur at the regulatory level (Wu et al. 1992; Oda et al. 2002). However, CRM degeneration, or “pseudo-enhancerization” has, to our knowledge, never been shown. Decreased TF binding levels could be a signal of degeneration, which could be caused by epigenetic or evolutionary changes at the sequence-level. Also, accelerated rate of evolution (relative to functional CRMs) could be a signal of relaxed selection on CRMs. Furthermore, in analogy to the pattern of K_a_/K_s_ in pseudogenes, there could be an excess of substitutions that decrease transcription factor binding site (TFBS) strength near pseudogenes compared to functional genes.

Here, we test these predictions by investigating whether relaxed constraint is identifiable in CRMs nearby the recently lost primate pseudogenes *Gulo* and *Uox*. We find evidence for decreased TF binding levels nearby both pseudogenes and lineage-specific relaxation of constraint in two putative regulatory regions near *Uox*.

We first investigated how the regulation of gene expression degenerates following pseudogenization of a coding region by focusing on the TF binding events of the liver transcription factors CEBPA, FOXA1, HNF4A and ONECUT1 (Ballester et al. 2014) nearby the two primate pseudogenes *Uox* and *Gulo*.

Although *Uox* is pseudogenized in the human-gibbon clade (Wu et al. 1992; Oda et al. 2002), TFs still bind nearby this locus (figure 1A). However, a clear difference is the loss of binding at the human promoter compared to the other four mammals (figure 1B). In contrast, there is no reduction in the number of ChIP-seq control (input) reads in this region (shown as gray histograms; figure 1) indicating that the absence of ChIP-seq signal is not due to lack of read mappability. Similar trends of lost binding can be seen in human and macaque at the *Gulo* locus as well (supplementary figure 1). Although these results support our prediction that regulatory degeneration would be accompanied by loss of transcription factor binding, evolutionary changes in mammalian transcription factor binding have been observed at the genome-wide scale (Schmidt et al. 2010; Ballester et al. 2014). We therefore sought to compare the patterns observed for pseudogenes to functional liver genes.

**Figure 1:**
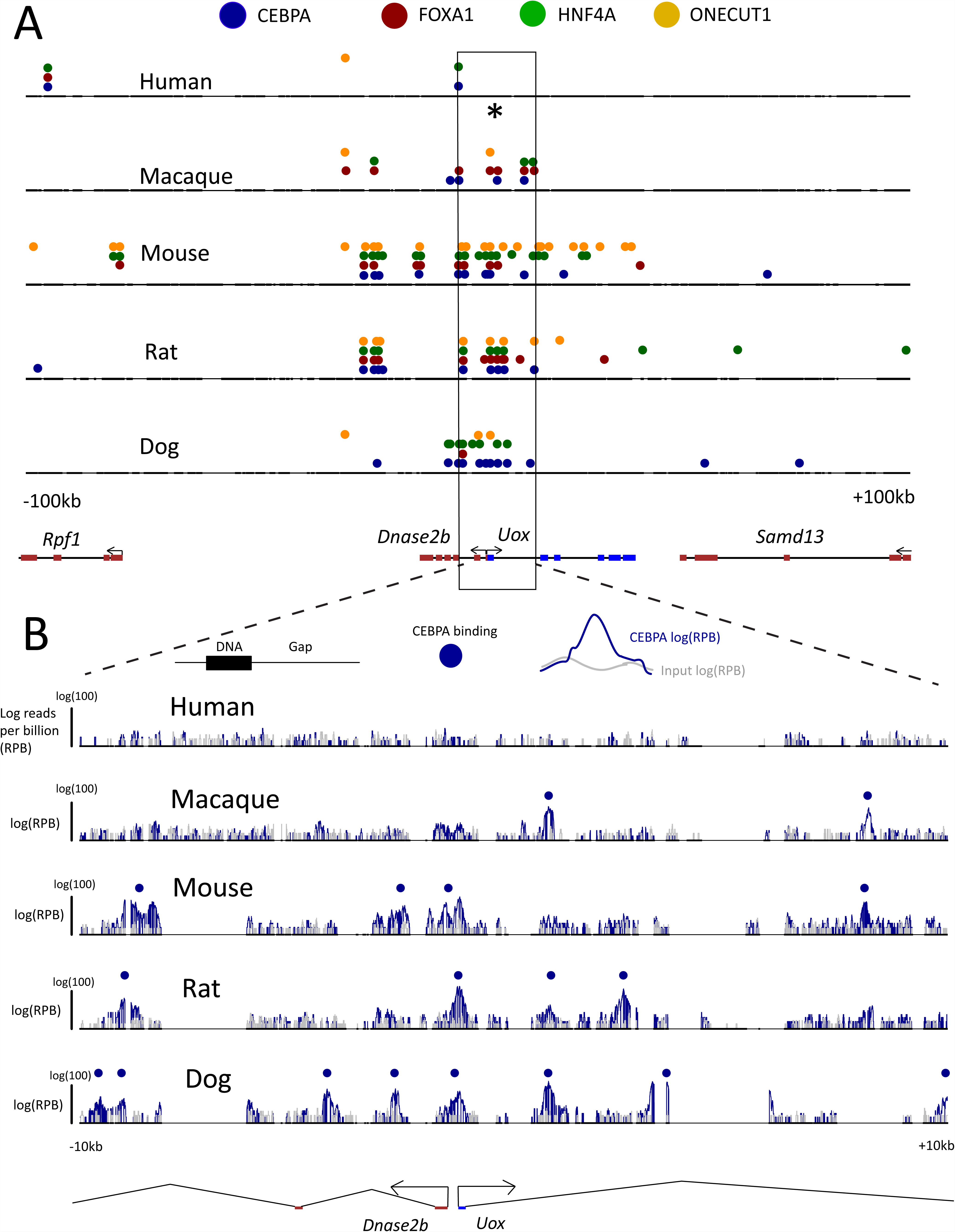
Multi-species alignment around the *Uox* locus in relative alignment coordinates. (A) An alignment corresponding to +/- 100kb around the human *Uox* TSS (chr1:84,763,514-84,963,514; negative strand). Thick black lines correspond to DNA for each species, while the thinner line indicates gaps. Different coloured circles illustrate the binding of the four TFs around this locus (as described in the legend above). * indicates the *Uox* promoter in humans, where there is marked loss in binding compared to the other species. (B) +/-10kb around the *Uox* TSS (indicated by box in panel A). The gray and blue histograms shows the distributions of input (a control, see materials and methods) and CEBPA reads per billion over the region in each species. The vertical black line at the leftmost side of each species name indicates log(100) reads per billion (RPB). Called CEBPA peaks in each species are indicated by blue circles. The *Uox* locus is coloured blue at the bottom of the alignment and nearby genes are shown in red.

We first confirmed that the loss of expression at pseudogene loci (as measured by RNA-seq) stands out relative to gene expression changes in liver expressed genes. Using a Brownian motion model (see materials and methods) we inferred gene expression in the rodent ancestor (because changes since the rodent ancestor can be estimated more reliably then changes since the primate ancestor) and then calculated the change in gene expression from this ancestor for these two pseudogenes and 1373 liver expressed genes. This large set of liver expressed genes was used because the inference of ancestral traits based upon only five extant species is noisy, so we wanted to use a large sample size to estimate the null distribution. The distributions of changes in gene expression along each lineage were converted into standard scores based upon the mean and standard deviation of the inferred changes in 1373 liver expressed genes (mean changes of ∼0 from the rodent ancestor to both human and macaque). The expression level decreases from the rodent ancestor to human for both pseudogenes relative to the 1373 liver genes (changes of -3.26 and -4.09 for *Gulo* and *Uox* respectively; Bonferroni corrected (BF corr.) combined *P* < 10^−6^; figure 2A). In contrast, while *Gulo* expression has significantly decreased in macaque (change: -3.42; BF corr. *P* < 10^−4^), *Uox* expression has not decreased more than expected by chance (change: -1.53; BF corr. *P* = 0.16; supplementary figure 2). This is expected because *Uox* is still functional and is only expressed moderately lower in macaque compared to other mammals (Oda et al. 2002).

**Figure 2:**
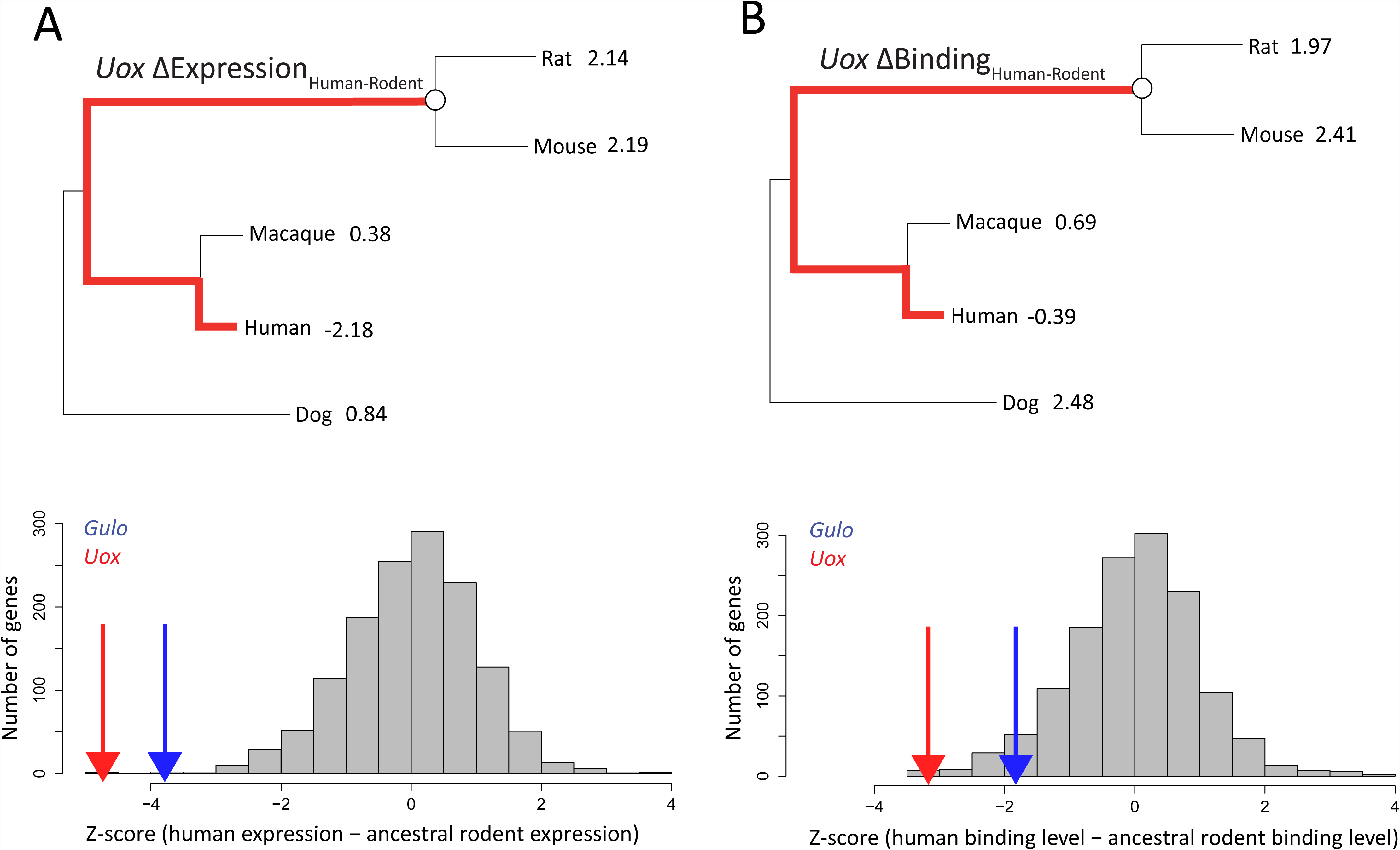
Both human gene expression and transcription factor binding level has decreased nearby primate pseudogenes. (A) The difference in expression level between the rodent ancestor and human. (B) The analogous difference in binding level between the rodent ancestor and human. Species trees show the inferred values of the quantitative traits. The white circles indicate the ancestral rodent and the red lines indicate the lineages being compared. Gray histograms correspond to the standard score of the distribution of changes for 1373 gene that are liver expressed in mouse, rat and dog with one-to-one orthologs across the five mammals. *Gulo* and *Uox* are indicated in terms of their standard scores by the blue and red lines respectively. In human, where both *Gulo* and *Uox* are pseudogenized, there is a significant decrease in both gene expression and binding level for the pseudogenes (combined *P* < 10^−4^).

We used a similar strategy to test whether the apparent decreased number of binding events in primate pseudogenes (figure 1) is statistically significant. We defined a quantitative measure of overall binding level that takes into account binding intensity, distance from a locus’ transcription start site (TSS) and the number of binding events (see materials and methods). This measure of binding level can be considered a quantitative trait for which we have observations in five species. If relaxation of selection has led to a loss of binding after pseudogenization, we predict that this quantitative trait will decrease in the human lineage. To test this, we computed the difference in human binding level for pseudogenes relative to the rodent ancestor, as above. We find a significant decrease in binding level along the human lineage for both pseudogenes relative to the 1373 liver genes (-1.46 and -2.46 for *Gulo* and *Uox* respectively, compared to a mean of -0.08 for liver expressed genes; BF corr. combined *P* < 10^−4^; figure 2B). The decrease in binding level in macaque is not significant for either *Gulo* or *Uox* (individual locus and combined *P* > 0.05; supplementary figure 2B).

We next tested for regulatory degeneration at the sequence level. Specifically, we tested for lineage-specific accelerated evolution in CRMs nearby *Uox*, which we expected in analogy to coding region degeneration, where the non-synonymous substitution rate (K_a_) along pseudogenized lineages accelerates relative to the synonymous (near-neutral) substitution rate (K_s_) (Torrents et al. 2002). A set of 23 mammalian liver-specific genes (see materials and methods) was used for these analyses since they should be enriched for functional binding events relative to the set of 1373 liver expressed genes used above. We first confirmed that the pseudogene-lineage-specific estimate of K_a_/K_s_ for *Uox* was an outlier among functional liver-specific genes (figure 3A). Along the pseudogenized lineage *Uox* has K_a_/K_s_=1.32 while the K_a_/K_s_=0.649 in macaque where *Uox* is still functional. The distribution of the differences in K_a_/K_s_ between the two primate sets for each gene clearly shows that this difference for *Uox* is an extreme outlier (figure 3A; 3.75 standard deviations). This is consistent with relaxed constraint on the *Uox* amino acid sequence.

**Figure 3:**
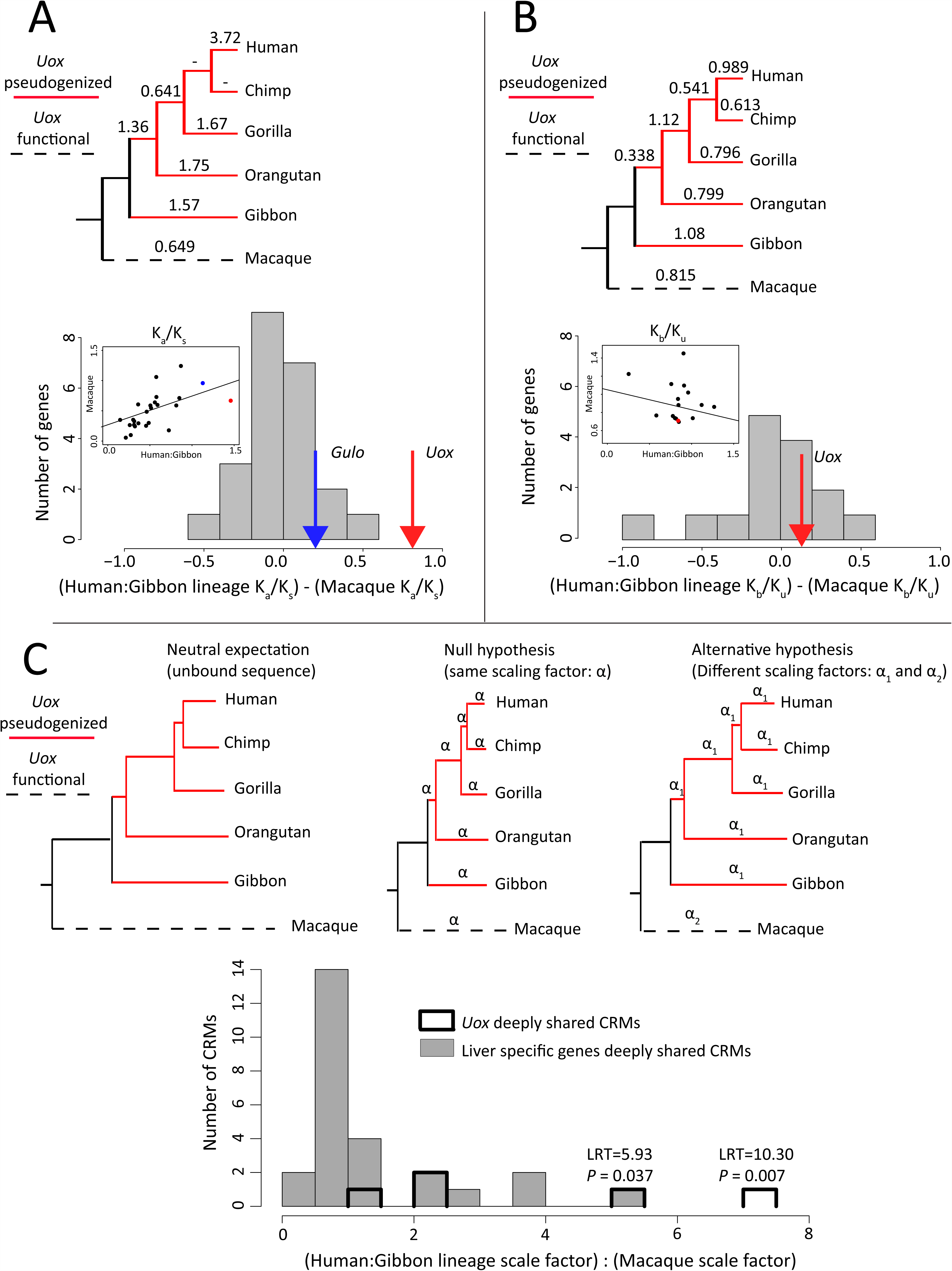
Tests for sequence degeneration. Red branches and internodes on phylogenies indicate pseudogenized lineages, while the dotted lines indicate that *Uox* is functional in macaque. (A) Comparisons of the non-synonymous substitution rate (K_a_), normalized by the synonymous (near-neutral) substitution rate (K_s_) along the human-gibbon lineage compared to macaque. The K_a_/K_s_ is indicated along each considered primate lineage (-indicates insufficient data). The grey histogram shows the background distribution for the difference in K_a_/K_s_ between the two sets of primates based upon the 23 liver-specific genes (each lineages’ Ka/K_s_ estimate is shown on the inset scatterplot). *Uox* is a clear outlier of this distribution (red arrow, 3.75 standard deviations away). (B) Comparisons of the bound sequence substitution rate (K_b_), normalized by the unbound sequence substitution rate (K_u_) along the human-gibbon lineage compared to macaque. The primate phylogeny indicates the K_b_/K_u_ along each considered primate lineage. The histogram and scatterplot are analogous to panel A, except they correspond to K_b_/K_u_ rather than K_a_/K_s_. *Uox* is not an outlier on this distribution of background differences (0.381 sd). (C) Likelihood ratio test of accelerated evolution in CRMs specifically along the human-gibbon lineage. This is a test of whether the scale factor for the rate of evolution in a CRM is the same or different between the human-gibbon lineage and macaque. Scale factors are relative to the branch lengths of a neutral reference phylogeny based upon unbound, flanking DNA. The three phylogenies represent this neutral reference phylogeny, the null (same scaling factors) model and the alternative (different scaling factor factor) models. Histograms indicate the distribution of scaling factors along the human-gibbon lineage relative to the macaque lineage for CRMs within 100kb of the liver-specific genes (grey) and *Uox* (white). 2/5 of the *Uox* CRMs have likelihood ratio test statistics (indicated on plot) that confidently reject the null hypothesis that the scaling factors are the same along the two lineages. There is only 1/27 CRM nearby liver-specific genes in the same range of scaling factors.

Analogous tests for human-gibbon accelerated substitution rates were used on putative regulatory sequences (CRMs) within 100kb of the *Uox* TSS. Here, regions bound by TFs were taken to be the sequences putatively under selection while unbound, non-coding flanking sequences were taken to be the neutral reference. We defined the ratio of substitutions in these regions to be K_b_/K_u_ (b=bound, u=unbound) in analogy to the K_a_/K_s_ ratio above (Hahn 2007). Peak sequences nearby the 23 liver-specific genes evolve more slowly than unbound flanking regions, both along the human-gibbon lineage (supplementary figure 3A; median of 0.0084 and 0.0072 sub. per site in human flank and peak sequences; Wilcoxon test *P* < 10^−6^) and in outgroup primates (supplementary figure 3B; median of 0.026 and 0.0245 in macaque flank and peak sequences respectively; Wilcoxon test BF corr. *P* = 0.02). This slower rate of evolution in peak sequences compared to flanks is consistent with purifying selection acting upon peak sequences and suggests that relaxed purifying selection should be detectable.

We therefore inferred the number of substitutions per site along the human-gibbon lineage and macaque branch in peaks nearby the liver-specific genes and plotted the difference in substitution rates inferred for each gene for all macaque peaks (figure 3B).The difference in rates for *Uox* falls within these distributions (0.381 sd). Therefore, there is no evidence for relaxed purifying selection on peak sequences nearby *Uox* compared to those nearby functional liver genes. We also investigated whether there is a difference in the pattern of changes in binding strength in TFBSs nearby these two gene-sets, but there was insufficient power at the single-gene level (Moses 2009; supplementary figure 4).

To investigate whether any particular CRMs show evidence for degeneration nearby *Uox* (putative “pseudo-enhancers”), we applied a likelihood ratio test (LRT) for accelerated evolution along the human-gibbon lineage. This tests whether a scale factor (relative to a neutral reference) for the rate of evolution within the two primate sets is significantly higher along the human-gibbon lineage than over the rest of the primate phylogeny (Cooper et al. 2005; Siepel et al. 2006; Figure 3C). We identified candidate highly conserved CRMs nearby *Uox* by requiring macaque binding events to be shared with at least two other non-human mammals. Within 100kb of the *Uox* transcription start sites there are five independent candidates for CRM degeneration based upon these criteria. We applied the LRT to test for accelerated evolution along the human-gibbon lineage. We found that 2/5 of these candidate degenerative CRMs are evolving significantly faster along the human-gibbon lineage (Figure 3C; FDR adjusted *P* < 0.05). These include multiple deeply shared binding events in the *Uox* promoter and a deeply shared ONECUT1 binding event in an intergenic region downstream of a neighbouring gene (see supplementary table 1 for all CRMs used in this analysis). This result is consistent with relaxed selection specifically along the human-gibbon lineage in the *Uox* promoter and a putative CRM nearby the *Uox* TSS.

We also tried to localize peaks that lost binding along pseudogenized lineages using the binding data alone. However, we were unable to do so because peaks with lost binding along pseudogenized lineages can be identified near functionally conserved liver genes. Thus, the multi-species ChIP-seq data is too variable to be used alone to identify candidate pseudo-enhancers. On the other hand, the quantitative decrease in TF binding (figure 2) suggests that at least some portion of TF binding events are preserved by purifying selection and that decreased binding levels reflect an increased proportion of TF binding events near pseudogenes being non-functional. Future work focusing on a larger set of pseudogenes, such as gene duplicates, could reveal whether decreased binding level is a general signature of regulatory degeneration. Additionally, although we did identify some candidate pseudo-enhancers using sequence-based analyses (figure 3C), the overall decrease in binding level could largely be driven by epigenetic changes instead.

The observation of accelerated evolution likely due to relaxed selection along the human-gibbon lineage in two regulatory elements (near the primate pseudogene *Uox*: the promoter and an intergenic CRM) is consistent with pseudo-enhancerization of regulatory elements in analogy to coding pseudogenization. In contrast, there was no signal of relaxed selection along the human-gibbon lineage when all bound sequences nearby *Uox* were combined (figure 3B). This result could suggest that a large proportion of bound sequences nearby gene-sets are non-functional (Cuvanovich et al. 2014) or under very weak selection. However, it is also possible that CRM degeneration is masked by the background turnover in TFBSs (Stergachis et al. 2014; Vierstra et al. 2014). This could be caused by decreased power to detect accelerated evolution due to the rapid gain and loss of functional TFBSs. Additionally, many regions of animal genomes have been identifiably bound by many different TFs simultaneously even though the known TFBSs of these TFs are not found (Moorman et al. 2006; The ENCODE Project Consortium 2012) so it is not clear that regulatory degeneration could be identified at the sequence level in these cases. Taken together our analysis suggests that although pseudo-enhancers may be identifiable in a small number of cases, decreased TF binding level over an entire locus could be a more reliable signature of regulatory degeneration.

## Materials and Methods

Methods are described in detail in the supplementary text, below is a brief summary.

We used publicly available ChIP-seq data produced from the livers of human, macaque, mouse, rat and dog for four liver-enriched TFs: CEBPA, FOXA1, HNF4A, ONECUT1 (Schmidt et al. 2010; Funnel et al. 2013; Steffalova et al. 2013; Ballester et al. 2014). Reads were mapped to each reference genome with Bowtie2 (Langmead and Salzberg 2012) with default parameters. Numbers of mapped reads and quality metrics are shown in supplementary table 2. Peak calling was performed with SWEMBL (v3.3.1; http://www.ebi.ac.uk/∼swilder/SWEMBL) with the parameter “-R 0.005”. Peaks were called as reproducible in both replicates if they overlapped by >=75% reciprocally (see supplementary table 3).

Based on a cut-off of >=5 FPKM within the livers of mouse, rat and dog (based upon previously published RNA-seq data, see supplementary text), a set of 1373 liver expressed genes that were one-to-one orthologs across the five mammals was constructed (“liver expressed genes”). Similarly, using Ensembl’s Expression Atlas (Cunningham et al. 2015), 23 one-to-one orthologs specific or overexpressed to each species’ liver of at least four of the mammals (and requiring both human and macaque to be included) were identified (“liver specific genes”).

The binding level for orthologs across species was based upon a previous approach (Wong et al. 2015). This approach combines the number of binding events, their binding intensity and their distance from the TSS into a single measure. Specifically, binding level (*a*_*is*_), where *i* is an orthologous gene in species *s* is given by:

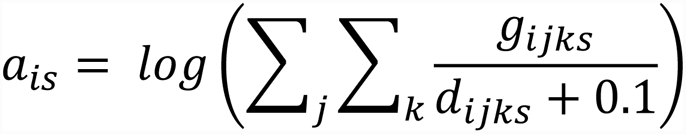

Where *k* is a peak within 100kb of the TSS of gene *i*, *g*_*ijks*_ is the intensity of peak *k* for TF *j* in species *s* and *d*_*ijks*_ is the distance (in bp) from the TSS to the summit of peak *k*. A pseudocount of 0.1 is added to the denominator to ensure this value is never zero. The mean and variance in binding level was calculated for all species and liver expressed genes (see transcriptome analysis). Since the signal-to-noise ratio differs greatly across ChIP-seq experiments, binding levels were converted into standard scores for each species so that they could be better compared. Maximum likelihood ancestral binding levels were reconstructed under a Brownian motion (BM) model in R using the “ace” function of the ape package (Paradis et al. 2004).

The complete 38 eutherian EPO alignment (Cunningham et al. 2015) was downloaded from the Ensembl FTP site (Release 74). Alignments were filtered if any of these five species were unalignable in this region. Neighbouring regions (+/- 2kb from peak ends) that did not intersect other peaks or exons were obtained. Shared peaks were called based upon overlapping summits (taken as the peak centre) within 150bp of each other. Primate sequences within the human-tarsier clade were parsed from these alignments. Ancestral sequences at each node in the primate phylogeny were reconstructed using the “prequel” program of PHAST (v1.3; Hubisz et al. 2011), which were used to call substitutions. Likelihood ratio tests were performed with the “phyloP” program of PHAST to test for accelerated evolution within the human-gibbon clade relative to the macaque branch.

## Acknowledgements

This work was supported by the Natural Sciences and Engineering Research Council of Canada.

